# Cell BLAST: Searching large-scale scRNA-seq databases via unbiased cell embedding

**DOI:** 10.1101/587360

**Authors:** Zhi-Jie Cao, Lin Wei, Shen Lu, De-Chang Yang, Ge Gao

## Abstract

An effective and efficient cell-querying method is critical for integrating existing scRNA-seq data and annotating new data. Herein, we present Cell BLAST, an accurate and robust cell-querying method. Powered by a well-curated reference database and a user-friendly Web server, Cell BLAST (http://cblast.gao-lab.org) provides a one-stop solution for real-world scRNA-seq cell querying and annotation.

## Main Text

Technological advances during the past decade have led to rapid accumulation of large-scale single-cell RNA sequencing (scRNA-seq) data. Analogous to biological sequence analysis^1^, identifying expression similarity to well-curated references via a cell-querying algorithm is becoming the first step of annotating newly sequenced cells. Tools have been developed to identify similar cells using approximate cosine distance^2^ or LSH Hamming distance^3, 4^ calculated from a subset of carefully selected genes. Such an intuitive approach is efficient, especially for large-scale data, but may suffer from nonbiological variation across datasets (batch effect^5, 6^). Meanwhile, multiple data harmonization methods have been proposed to remove such confounding factors during alignment, for example, via warping canonical correlation vectors^7^ or matching mutual nearest neighbors across batches^6^. While these methods can be applied to align multiple reference datasets, computation-intensive realignment is required to map query cells to the (pre-)aligned reference data space.

To address these challenges, we introduce a new customized deep generative model together with a novel cell-to-cell similarity metric specifically designed for cell querying (**Fig. 1a**, **Method**). Differing from canonical variational autoencoder (VAE) models^8-11^, adversarial batch alignment is applied to correct batch effect during low-dimensional embedding of reference datasets. Such a design also enables a special “online tuning” mode that can handle batch effect between query and reference data when necessary. Moreover, by exploiting the model’s universal approximator posterior to model uncertainty in latent space, we implement a distribution-based metric to measure cell-to-cell similarity. Finally, we also provide a well-curated multispecies single-cell transcriptomics database (ACA) and an easy-to-use Web interface for convenient exploratory analysis.

**Figure. 1.**
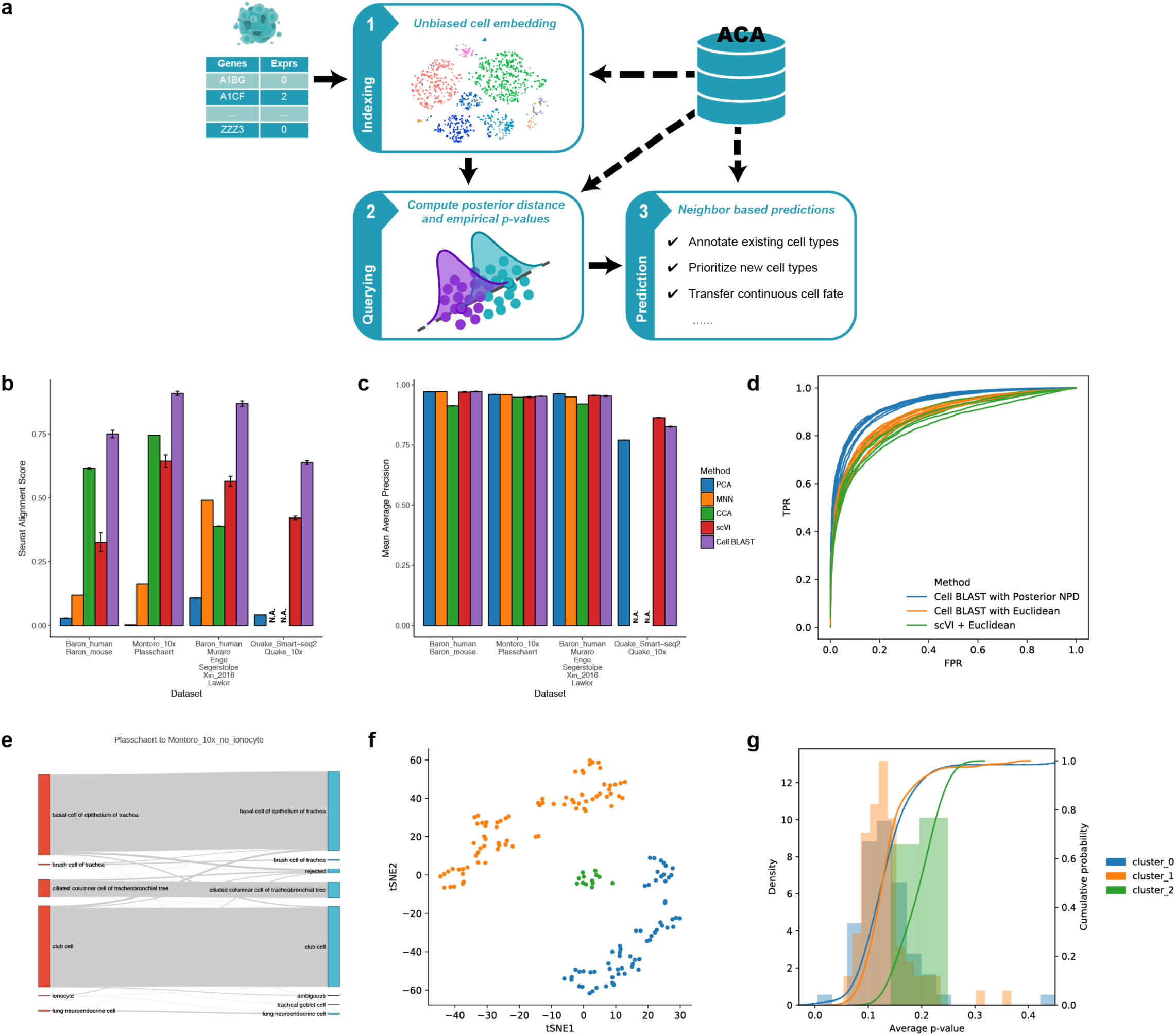
Cell BLAST benchmarking and application to trachea datasets. (**a**) Overall Cell BLAST workflow. (**b**) Extent of dataset mixing after batch effect correction in four groups of datasets, quantified by the Seurat alignment score. A high Seurat alignment score indicates that local neighborhoods consist of cells from different datasets uniformly rather than from the same dataset only. Error bars indicate mean ± s.d. Methods that did not finish under the 2-hour time limit are marked as N.A. (**c**) Cell type resolution after batch effect correction, quantified by cell type mean average precision (MAP). MAP can be thought of as a generalization to nearest neighbor accuracy, with larger values indicating higher cell type resolution, thus more suitable for cell querying. Error bars indicate mean ± s.d. Methods that did not finish under the 2-hour time limit are marked as N.A. (**d**) ROC curve of different distance metrics in discriminating cell pairs with the same cell type from cell pairs with different cell types. (**e**) Sankey plot comparing Cell BLAST predictions and original cell type annotations for the “Plasschaert” dataset. (**f**) t-SNE visualization of Cell BLAST-rejected cells, colored by unsupervised clustering. (**g**) Average p-value distribution of each cluster in (**f**).

To assess our model’s capability of capturing biological similarity in the low-dimensional latent space, we first benchmarked against several popular dimension reduction tools^8, 12, 13^ using real-world data (**Supplementary Table 1**) and found that our model is overall among the best performing methods (**Supplementary Fig. 1-2**). We further compared batch effect correction performance using combinations of multiple datasets with overlapping cell types profiled (**Supplementary Table 1**). Our model achieves significantly better dataset mixing (**Fig. 1b**) while maintaining comparable cell type resolution (**Fig. 1c**). Latent space visualization also demonstrates that our model can effectively remove batch effect for multiple datasets with a considerable difference in cell type distribution (**Supplementary Fig. 3**). Notably, we found that the correction of inter-dataset batch effect does not automatically generalize to that within each dataset, which is most evident in the pancreatic datasets (**Supplementary Fig. 3c-d**, **Supplementary Fig. 4a-c**). For such complex scenarios, our model is effective in removing multiple levels of batch effect simultaneously (**Supplementary Fig. 4d-h**).

While the unbiased latent space embedding derived by the nonlinear deep neural network effectively removes confounding factors, the network’s random components and nonconvex optimization procedure also lead to serious challenges, especially false-positive hits when cells outside reference types are provided as query. Thus, we propose a novel posterior distribution-based cell-to-cell similarity metric in the latent space, which we term “normalized projection distance” (NPD). Distance metric ROC analysis (**Method**) shows that our posterior NPD metric is more accurate and robust than Euclidean distance which is commonly used in other neural network-based embedding tools (**Fig. 1d**, **Supplementary Fig. 4k**). Additionally, we exploit the stability of query-hit distance across multiple models to improve specificity (**Method**, **Supplementary Fig. 4l**). An empirical p-value is computed for each query hit as a measure of “confidence” by comparing the posterior distance to the empirical NULL distribution obtained from randomly selected pairs of cells in the queried data.

The high specificity of Cell BLAST is especially important for discovering novel cell types. Two recent studies (“Montoro”^14^ and “Plasschaert”^15^) independently reported a rare tracheal cell type named pulmonary ionocyte. We artificially removed ionocytes from the “Montoro” dataset and used it as a reference to annotate query cells from the “Plasschaert” dataset. In addition to accurately annotating 95.9% of query cells, Cell BLAST correctly rejected 12 of 19 “Plasschaert” ionocytes (**Fig. 1e**). Moreover, it highlights the existence of a putative novel cell type as a well-defined cluster with large p-values among all 156 rejected cells (**Fig. 1f-g**). Further examination shows that this cluster actually corresponds to ionocytes (**Supplementary Fig. 6a**; also see **Supplementary Fig. 5** for more detailed analysis on the remaining 7 ionocytes). By contrast, scmap-cell^2^ only rejected 7 “Plasschaert” ionocytes despite the higher overall rejection number of 401 (i.e., more false negatives; **Supplementary Fig. 6b-e**).

We further systematically compared the performance of query-based cell typing with scmap-cell^2^ and CellFishing.jl^4^ (**Method**) using four groups of datasets, each including both positive control and negative control queries (first 4 groups in **Supplementary Table 2**). Of interest, while Cell BLAST shows superior performance than scmap-cell and CellFishing.jl under the default setting (**Supplementary Fig. 7a-c**, **8-10**), detailed ROC analysis reveals that the performance of scmap-cell could be further improved to a level comparable to Cell BLAST by employing higher thresholds, while ROC and optimal thresholds of CellFishing.jl show large variation across different datasets (**Supplementary Fig. 7d**). Cell BLAST presents the most robust performance with a default threshold (p-value < 0.05) across different datasets, which will significantly benefit real-world application. Additionally, we assessed their scalability using reference data varying from 1,000 to 1,000,000 cells. Both Cell BLAST and CellFishing.jl scale well with increasing reference size, while scmap-cell’s querying time rises dramatically for larger reference datasets with more than 10,000 cells (**Supplementary Fig. 7e**).

Moreover, our deep generative model combined with posterior-based latent-space similarity metric enables Cell BLAST to model the continuous spectrum of cell states accurately. We demonstrate this using a study profiling mouse hematopoietic progenitor cells (“Tusi”^16^) in which computationally inferred cell fate distributions are available. For the purpose of evaluation, cell fate distributions inferred by the authors are recognized as ground truth. We selected cells from one sequencing run as query and the other as reference to test whether we can accurately transfer continuous cell fate between experimental batches (**Fig. 2a-b**). Jensen-Shannon divergence between predicted cell fate distributions and ground truth shows that our prediction is again more accurate than scmap (**Fig. 2c**).

**Figure. 2.**
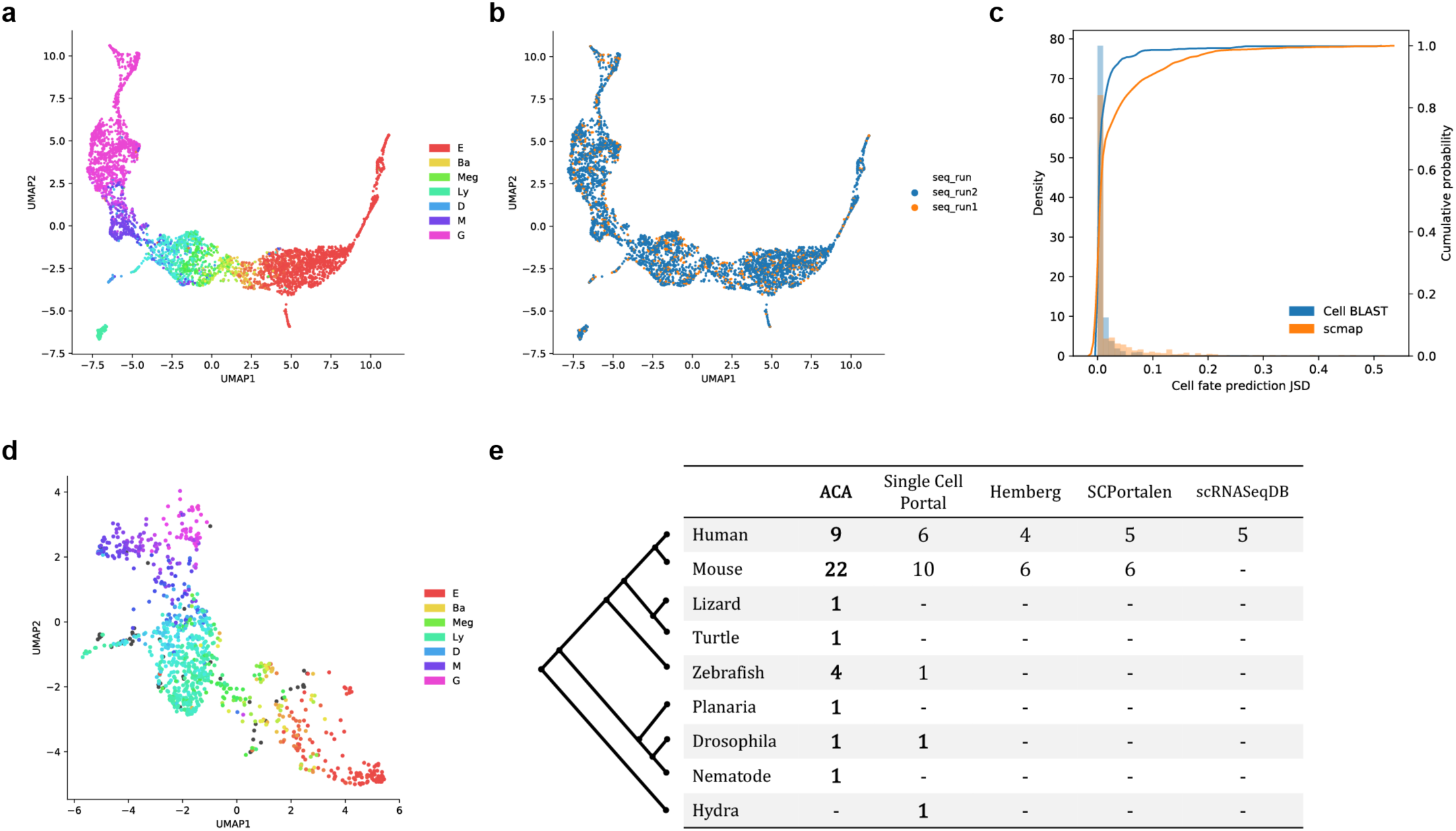
Application to hematopoietic progenitor datasets. (**a**, **b**) UMAP visualization of latent space learned on the “Tusi” dataset, colored by sequencing run (**a**) and cell fate (**b**). The model is trained solely on cells from run 2 and used to project cells from run 1. Each of the seven terminal cell fates (E, erythroid; Ba, basophilic or mast; Meg, megakaryocytic; Ly, lymphocytic; D, dendritic; M, monocytic; G, granulocytic neutrophil) is assigned a distinct color. The color of each single cell is then determined by the linear combination of these seven colors in hue space, weighed by the cell fate distribution among these terminal fates. (**c**) Distribution of Jensen-Shannon divergence between predicted cell fate distributions and author-provided “ground truth“. (**d**) UMAP visualization of the “Velten” dataset, colored by Cell BLAST-predicted cell fates. (**e**) Number of organs covered in each species for different single-cell transcriptomics databases, including the Single Cell Portal (https://portals.broadinstitute.org/single_cell), Hemberg collection^2^, SCPortalen^19^, and scRNASeqDB^20^.

Besides batch effect among different reference datasets, *bona fide* biological similarity could also be confounded by large, undesirable bias between query and reference data. Exploiting the dedicated adversarial batch alignment, we implemented a particular “online tuning” mode to handle such an often-neglected confounding factor. Briefly, the combination of reference and query data is used to fine-tune the existing reference-based model, with the query-reference batch effect added as an additional component to be removed by adversarial batch alignment (**Method**). Using this strategy, we successfully transferred cell fate from the above “Tusi” dataset to an independent human hematopoietic progenitor dataset (“Velten”^17^) (**Fig. 2d**). The expression of known cell lineage markers validates the rationality of transferred cell fates (**Supplementary Fig. 11a-f**). By contrast, scmap-cell incorrectly assigned most cells to monocyte and granulocyte lineages (**Supplementary Fig. 11g**). As another example, we applied “online tuning” to *Tabula Muris*^18^ spleen data, which exhibit significant batch effect between 10x- and Smart-seq2-processed cells. The ROC of Cell BLAST improved significantly after “online tuning”, achieving high specificity, sensitivity and Cohen’s κ (a measure of prediction accuracy corrected for chance, see **Methods** for more details)^2^ at the default cutoff (**Supplementary Fig. 11h**, last group in **Supplementary Table 2**).

A comprehensive and well-curated reference database is crucial for the practical application of Cell BLAST. Based on public scRNA-seq datasets, we curated ACA, a high-quality reference database. With 986,305 cells in total, ACA currently covers 27 distinct organs across 8 species, offering the most comprehensive compendium for diverse species and organs (**Fig. 2e**, **Supplementary Fig. 12a-b**, **Supplementary Table 3**). To ensure a unified and high-resolution cell type description, all records in ACA are collected and annotated using a standard procedure (**Method**), with 98.9% of datasets manually curated with Cell Ontology, a structured controlled vocabulary for cell types. We trained our model on all ACA datasets. Notably, we found that the model works well in most cases with minimal hyperparameter tuning (latent space visualizations, self-projection coverage and accuracy available on our website, **Supplementary Fig. 12e**).

A user-friendly Web server is publicly accessible at http://cblast.gao-lab.org, with all curated datasets and pretrained models available. Based on the wealth of resources, our website provides “off-the-shelf” querying service. Users can obtain querying hits and visualize cell type predictions with minimal effort (**Supplementary Fig. 12c-d**). For advanced users, a well-documented Python package implementing the Cell BLAST toolkit is also available, which enables model training on custom references and diverse downstream analyses.

By explicitly modeling multilevel batch effect as well as uncertainty in cell-to-cell similarity estimation, Cell BLAST is an accurate and robust querying algorithm for heterogeneous single-cell transcriptome datasets. In combination with a comprehensive, well-annotated database and an easy-to-use Web interface, Cell BLAST provides a one-stop solution for both bench biologists and bioinformaticians.

### Software availability

The full package of Cell BLAST is available at http://cblast.gao-lab.org. Code necessary to reproduce results in the paper is deposited at https://github.com/gao-lab/Cell_BLAST and https://github.com/gao-lab/Cell_BLAST-notebooks.

## Acknowledgments

The authors thank Drs. Zemin Zhang, Cheng Li, Letian Tao, Jian Lu and Liping Wei at Peking University for their helpful comments and suggestions during the study.

This work was supported by funds from the National Key Research and Development Program (2016YFC0901603), the China 863 Program (2015AA020108), as well as the State Key Laboratory of Protein and Plant Gene Research and the Beijing Advanced Innovation Center for Genomics (ICG) at Peking University. The research of G.G. was supported in part by the National Program for Support of Top-notch Young Professionals.

Part of the analysis was performed on the Computing Platform of the Center for Life Sciences of Peking University and supported by the High-performance Computing Platform of Peking University.

## Author contributions

G.G. conceived the study and supervised the research; Z.J.C. and L.W. contributed to the computational framework and data curation; S. L., Z.J.C., and D.C.Y designed, implemented and deployed the website; Z.J.C. and G.G. wrote the manuscript with comments and inputs from all coauthors.

## Methods

### The deep generative model

The model we used is based on the adversarial autoencoder (AAE)^21^. Below, we denote the gene expression profile of a cell as ***x*** ∈ ℝ^*G*^, where *G* is the number of genes. The data generative process is modeled by a continuous latent variable ***z*** ∈ ℝ^*D*^(*D* ≪ *G*) with standard Gaussian prior ***z*** ∼ *N*(**0**, *I*_*D*_) which models continuous cell states, as well as a one-hot latent variable ***c*** ∈ {0,1}^*k*^, ***c***^*T*^***c*** = 1 with categorical prior ***c*** ∼ *Cat*(*K*) which aims to model cell type clusters. A unified latent vector is then determined by ***l*** = ***z*** + *H**c***, where *H* ∈ ℝ^D×K^. A neural network (decoder, denoted by Dec below) maps the cell embedding vector ***l*** to two parameters of the negative binomial distribution ***μ**, **θ*** = Dec(***l***) that models the distribution of expression profile ***x***:

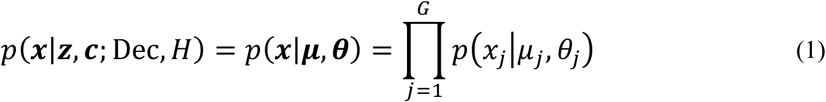

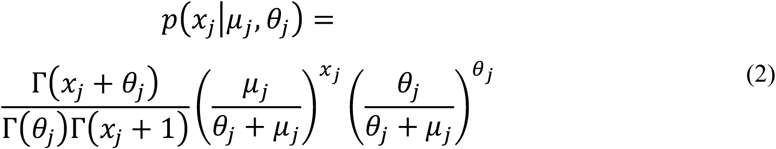

where ***μ*** and ***θ*** are the mean and dispersion of the negative binomial distribution, respectively. Theoretically, the negative binomial model should be fitted on raw count data^8, 13, 22^. However, for the purpose of cell querying, datasets have to be normalized to minimize the influence of capture efficiency and sequencing depth. We empirically found that, using normalized data, the negative binomial model still produced better results than alternative distributions like the log-normal distribution. To prevent numerical instability during training caused by normalization that breaks the mean-variance relationship of the negative binomial model, we additionally included the variance of the dispersion parameter as a regularization term.

Training objectives for the adversarial autoencoder are:

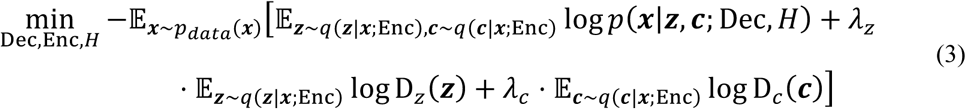

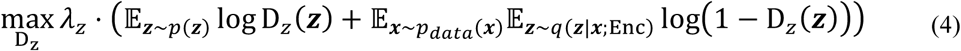

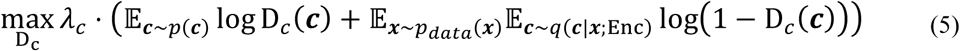

*q*(***z***|***x***; Enc) and *q*(***c***|*x*; Enc) are “universal approximator posteriors” parameterized by another neural network (encoder, denoted by Enc). Expectations with regard to *q*(***z***|***x***; Enc) and *q*(***c***|***x***; Enc) are approximated by sampling ***x*****’** ∼ *poisson*(***x***) and feeding to the deterministic encoder network. The choice of Poisson noise is arbitrary as the encoder learns to map this arbitrary noise distribution to an appropriate posterior distribution during training. D_z_ and D_c_ are discriminator networks for ***z*** and ***c***, respectively, which output the probability that a latent sample is from the prior rather than from the posterior. Effectively, adversarial training between the encoder (Enc) and discriminators (D_*z*_ and D_*c*_) drives the encoder output to match prior distributions of latent variables *p*(***z***) and *p*(***c***). *λ*_*z*_ and *λ*_*c*_ are hyperparameters that control prior matching strength. The model is much easier and more stable to train than canonical GANs because of the low dimensionality and simple distribution of ***z*** and ***c***.

At convergence, the encoder learns to map the data distribution to latent variables that follow their respective prior distributions, and the decoder learns to map latent variables from prior distributions back to the data distribution. The key element we use for cell querying is vector ***l*** on the decoding path because it defines a unified latent space in which biological similarities are well captured. The model also works if no categorical latent variable is used, in which case ***l*** = ***z*** directly.

Some architectural designs are learned from scVI^8^, including logarithm transformation before encoder input, and softmax output scaled by the library size when computing ***μ***. Stochastic gradient descent with minibatches is applied to optimize the loss functions. Specifically, we use the “RMSProp” optimization algorithm with no momentum term to ensure stability of adversarial training. The model is implemented using the Tensorflow^23^ Python library.

### Adversarial batch alignment

As a natural extension to the prior matching adversarial training strategy described in the previous section, and following recent work in domain adaptation^24-26^, we propose the adversarial batch alignment strategy to align the latent space distribution of different batches. We denote the batch membership of each cell as ***b*** ∈ {0,1}^*B*^, ***b***^*T*^***b*** = 1. The distribution *p*(***b***) is categorical:

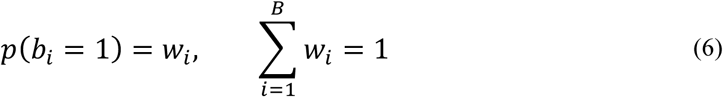

Adversarial batch alignment introduces an additional loss:

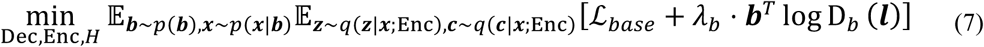

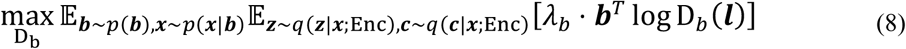

*ℒ*_*base*_ denotes the loss function in (3). D_*b*_ is a multiclass batch discriminator network that outputs the probability distribution of batch membership based on the embedding vector ***l***. *λ*_*b*_ is a hyperparameter controlling batch alignment strength. Additionally, the generative distribution is extended to condition on ***b*** as well:

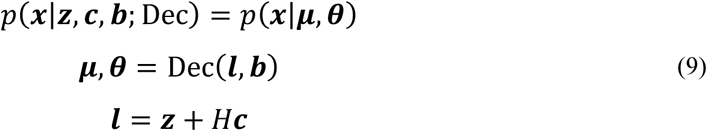

Below, we focus on batch alignment and discard the first *ℒ*_*base*_ term and scaling parameter *λ*_*b*_. We extend the derivation in the original GAN paper^27^ to show that adversarial batch alignment converges when embedding space distributions of different batches are aligned. To simplify notation, we fuse the data distribution and encoder transformation and replace the minimization over encoder to minimization over batch-embedding distributions:

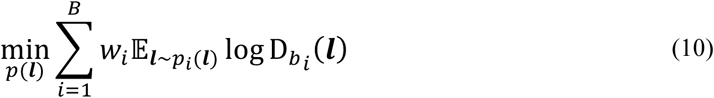

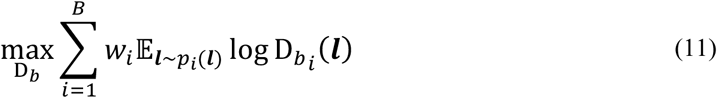

Here 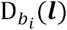 denotes the *i*^th^ dimension of the discriminator output, i.e., the probability that the discriminator “thinks” a cell is from the *i*^th^ batch. D_*b*_ is assumed to have sufficient capacity, which is generally reasonable in the case of neural networks. The global optimum of (11) is reached when D_*b*_ outputs optimal batch membership distribution at every ***l***:

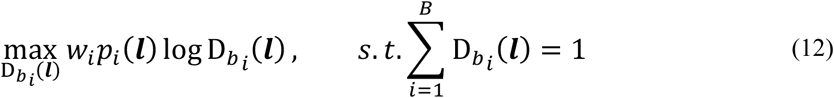

The solution to the above maximization is given by:

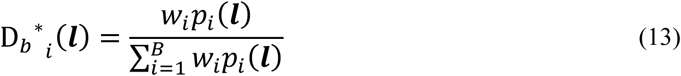

Substituting D_*b*_ *(*l*) back into (10), we obtain:

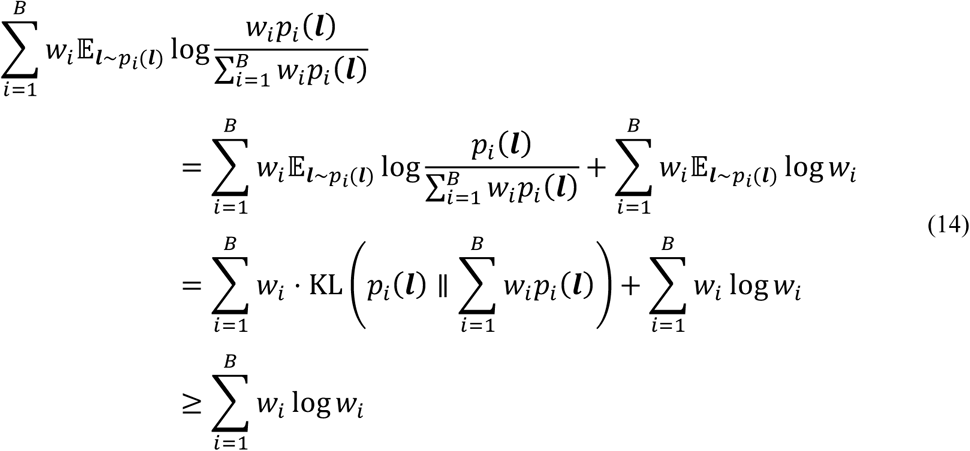

Thus, 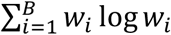 is the global minimum, reached if and only if *p*_*i*_(*l*) = *p*_*j*_(*l*), ∀*i, j*. The minimization of (10) is equivalent to minimizing a form of generalized Jensen-Shannon divergence among multiple batch-embedding distributions.

Note that in practice, model training balances between *ℒ*_*base*_ and pure batch alignment. Aligning cells of the same type induces a minimal cost in *ℒ*_*base*_, while improperly aligning cells of different types could cause *ℒ*_*base*_ to rise dramatically. During training, the gradient from both batch discriminators and decoder provide fine-grain guidance to align different batches, leading to better results than “hand-crafted” alignment strategies like CCA^7^ and MNN^6^. Empirically, given proper values for *λ*_*b*_, the adversarial approach correctly handles difference in cell type distribution among batches. If multiple levels of batch effect exist, e.g., within-dataset and cross-dataset, we use an independent batch discriminator for each component, providing extra flexibility.

### Data preprocessing for benchmarks

Most informative genes were selected using the Seurat^7^ function “FindVariableGenes”. We set the argument “binning.method” to “equal_frequency” and left other arguments as default. If within-dataset batch effect exists, genes are selected independently for each batch and then pooled together. By default, a gene is retained if it is selected in at least 50% of batches. Downstream benchmarks were all performed using this gene set, except for scmap and CellFishing.jl, which provide their own gene selection method. GNU parallel^28^ was used to parallelize and manage jobs throughout the benchmarking and data processing pipeline.

### Benchmarking dimension reduction

PCA was performed using the R package irlba^29^ (v2.3.2). ZIFA^12^ was downloaded from its Github repository, and hard coded random seeds were removed to reveal actual stability. ZINB-WaVE^13^ (v1.0.0) was performed using the R package zinbwave. scVI^8^ (v0.2.3) was downloaded from its Github repository, and minor changes were made to the original code to address PyTorch^30^ compatibility issues. Our modified versions of ZIFA and scVI are available upon request.

For PCA and ZIFA, data were logarithm transformed after normalization and adding a pseudocount of 1. Hyperparameters of all methods above were left as default. For our model, we used the same set of hyperparameters throughout all benchmarks. *λ*_*z*_ and *λ*_*c*_ were both set to 0.001. All neural networks (encoder, decoder and discriminators) used a single layer of 128 hidden units. Learning rate of the RMSProp optimizer is set to 0.001, and minibatches of size 128 were used. For comparability, the target dimensionality of each method was set to 10. All benchmarked methods were repeated multiple times with different random seeds. 4 random seeds were used for PCA, ZIFA and ZINB-WaVE, while 16 random seeds were used for scVI and our model, since neural network-based models are typically considered less stable. Run time was limited to 2 hours, after which the jobs were terminated.

Cell type nearest neighbor mean average precision (MAP) was computed with K nearest neighbors of each cell based on low-dimensional space Euclidean distance. If we denote the cell type of a cell as *y*, and the cell types of its ordered nearest neighbors as *y*_1_, *y*_2_, & *y*_*k*_. The average precision (AP) for that cell is defined as:

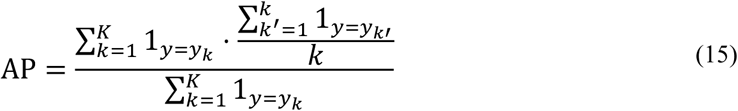

Mean average precision is then given by:

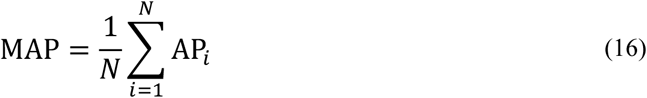

Note that when *K* = 1, MAP reduces to the nearest neighbor accuracy. We set *K* to 1% of the total cell number throughout all benchmarks.

### Benchmarking batch effect correction

We merged multiple datasets according to shared gene names. If datasets to be merged are from different species, Ensembl ortholog^31^ information was used to map genes to ortholog groups before merging. To obtain informative genes in merged datasets, we take the union of informative genes from each dataset, and then intersect the union with the intersection of detected genes from each dataset.

CCA^7^ and MNN^6^ alignments were performed using the R packages Seurat^7^ (v2.3.3) and scran^32^ (v1.6.9), respectively. Hard-coded random seeds in Seurat were removed to reveal actual stability. The modified version of Seurat is available upon request. For comparability, we evaluated cell type resolution and batch mixing in a 10-dimensional latent space. For MNN alignment, we set the argument “cos.norm.out” to false and left other arguments as default. PCA was applied to reduce the dimensionality to 10 after obtaining the MNN-corrected expression matrix. For CCA alignment, we used the first 10 canonical correlation vectors. Run time was limited to 2 hours, after which the jobs were terminated. Seurat alignment score was computed exactly as described in the CCA alignment paper^7^. For our own model, we consistently used *λ*_*b*_ = 0.01, and all other hyperparameters remain the same as in dimension reduction benchmarks. 4 random seeds were used for PCA, CCA and MNN, while 16 random seeds were used for scVI and our model, since neural network-based models are typically considered less stable.

### Cell querying based on posterior distributions

We evaluated cell-to-cell similarity based on the posterior distribution distance. Similar to the training phase, we obtained samples from the “universal approximator posterior” by sampling ***x*****’** ∼ *Poisson*(***x***) and feeding to the encoder network. To obtain a robust estimation of the distribution distance with a small number of posterior samples, we project the posterior samples of two cells onto the line connecting their posterior point estimates in the latent space and use the projected scalar distribution distance to approximate the true distribution distance. Wasserstein distance is computed on normalized projections to account for nonuniform density across the embedding space:

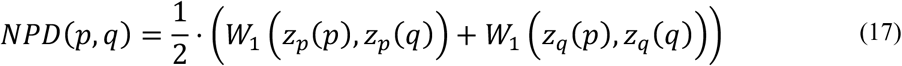

Where

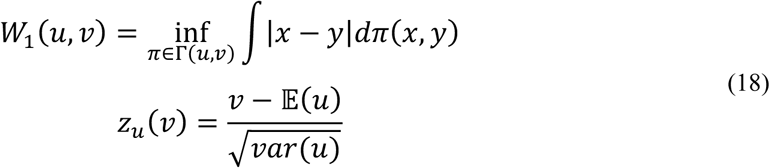

We term this distance metric normalized projection distance (NPD). By default, 50 samples from the posterior are used to compute NPD, which produces sufficiently accurate results (**Supplementary Fig. 4i-j**). The definition of posterior NPD does not imply an efficient nearest neighbor searching algorithm. To increase speed, we first use Euclidean distance-based nearest neighbor searching, which is highly efficient in the low-dimensional latent space, and then compute posterior distances only for these nearest neighbors. The empirical distribution of posterior NPD for a dataset is obtained by computing posterior NPD on randomly selected pairs of cells in the reference dataset. Empirical p-values of query hits are computed by comparing the posterior NPD of a query hit to this empirical distribution. We note that even with the querying strategy described above, querying with single models still occasionally leads to many false-positive hits when cell types on which the model has not been trained are provided as query. This is because embeddings of such untrained cell types are mostly random, and they could localize close to reference cells by chance. We reason that embedding randomness of untrained cell types could be utilized to identify and correctly reject them. Practically, we train multiple models with different starting points (as determined by random seeds) and compute query hit significance for each model. A query hit is considered significant only if it is consistently significant across multiple models. To acquire predictions based on significant hits, we use majority voting for discrete variables, e.g., cell type, or averaging for continuous variables, e.g., cell fate distribution.

### Distance metric ROC analysis

Our model and scVI^8^ were fitted on reference datasets and applied to positive and negative control query datasets in the pancreas group of **Supplementary Table 2**. We then randomly selected 10,000 query-reference cell pairs. A query-reference pair is defined as “positive” if the query cell and reference cell are of the same cell type, and “negative” otherwise. Each benchmarked similarity metric was then computed on all sampled query-reference pairs and used as predictors for “positive”/“negative” pairs. AUROC values were computed for each benchmarked similarity metric. In addition to the Euclidean distance, we also computed posterior distribution distances for scVI (**Supplementary Fig. 4k**). NPD was computed as described in (17), based on samples from the posterior Gaussian. JSD was computed in the original latent space without projection.

### Benchmarking query-based cell typing

Cell ontology annotations in ACA were used as ground truth. Cells without cell ontology annotations were excluded in the analysis. For each querying method, cell type predictions for query cells were obtained based on query hits with a minimal similarity cutoff, i.e., query cells with no significant hits are rejected, while cells not rejected are further assigned cell type predictions. Sensitivity, specificity and Cohen’s κ are computed as follows:

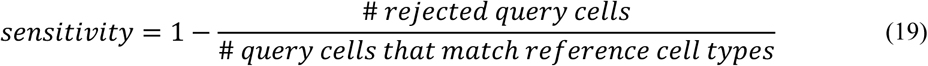

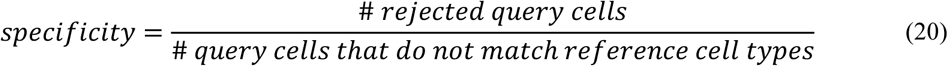

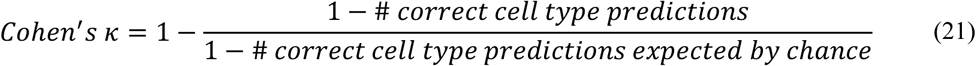

Predictions are considered correct if they exactly match the ground truth, i.e., no flexibility based on cell type similarity. This prevents unnecessary bias introduced in the selection of cell type similarity measure. Cells were inversely weighed by the size of the corresponding dataset when computing average sensitivity, specificity and Cohen’s κ. AUROC was computed using linear interpolation. For scmap^2^, we varied the minimal cosine similarity requirement for nearest neighbors. For Cell BLAST, we varied the maximal p-value cutoff used in filtering hits. For CellFishing.jl^4^, the original implementation does not include a dedicated cell type prediction function, so we used the same strategy as that for our own method (majority voting after distance filtering) to acquire final predictions, in which we varied the Hamming distance cutoff used in distance filtering. Finally, 4 random seeds were tested for each cutoff and each method to reflect stability. Several other cell querying tools (CellAtlasSearch^3^, scQuery^33^, scMCA^34^) were not included in our benchmark because they do not support custom reference datasets.

### Benchmarking querying speed

To evaluate the scalability of querying methods, we constructed reference datasets of varying sizes by subsampling from the 1M mouse brain dataset^35^. For query data, the “Marques” dataset^36^ was used. Benchmarking was performed on a workstation with 40 CPU cores, 100GB RAM and GeForce GTX 1080Ti GPU. For all methods, only the querying time was recorded, not including the time consumed to build reference indices.

### Application to trachea datasets

We first removed cells labeled as “ionocytes” in the “Montoro_10x”^14^ dataset and used “FindVariableGenes” from Seurat to select informative genes in the remaining cells. Four models with different starting points were trained on the tampered “Montoro_10x” dataset. We used a cutoff of empirical p-value > 0.1 to reject query cells from the “Plasschaert”^15^ dataset as potential novel cell types. We clustered rejected cells using spectral clustering (Scikit-learn^37^ v0.20.1) after applying t-SNE^38^ to latent space coordinates. The average p-value for a query cell was computed as the geometric mean of p-values across all hits.

### Online tuning

When significant batch effect exists between reference and query, we support further aligning query data with the reference data in an online-learning manner. All components in the pretrained model, including the encoder, decoder, prior discriminators and batch discriminators, are retained. The reference-query batch effect is added as an extra component to be removed using adversarial batch alignment. Specifically, a new discriminator dedicated to the reference-query batch effect is added, and the decoder is expanded to accept an extra one-hot indicator for reference and query. The expanded model is then fine-tuned using the combination of reference and query data. Two precautions are taken to prevent a decrease in specificity caused by over-alignment. First, adversarial alignment loss is constrained to cells that have mutual nearest neighbors^6^ between reference and query data in each SGD minibatch. Second, we penalize the deviation of tuned model weights from the original weights.

### Application to hematopoietic progenitor datasets

For the within-“Tusi”^16^ query, we trained four models using only cells from sequencing run 2, and cells from sequencing run 1 were used as query cells. PBA inferred cell fate distributions provided by the authors, which are 7-dimensional categorical distributions across 7 terminal cell fates, were used as the ground truth. We took the average cell fate distributions of significant querying hits (p-value < 0.05) as predictions for query cells. Regarding scmap-cell, we filtered nearest neighbors according to a default cosine similarity cutoff of 0.5. Jensen-Shannon divergence (JSD) between true and predicted cell fate distributions was computed as below:

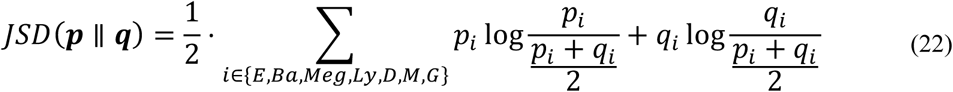

For cross-species querying between “Tusi” and “Velten”^17^, we mapped both mouse and human genes to ortholog groups. Online tuning with 200 epochs was used to increase sensitivity and accuracy. Latent space visualization was performed using UMAP^39, 40^.

### ACA database construction

We searched Gene Expression Omnibus (GEO)^41^ using the following search term:

~~~
(
  "expression profiling by high throughput sequencing"[DataSet Type] OR
  "expression profiling by high throughput sequencing"[Filter] OR
  "high throughput sequencing"[Platform Technology Type]
) AND
"gse"[Entry Type] AND
(
"single cell"[Title] OR
"single-cell"[Title]
) AND
("2013"[Publication Date]: "3000"[Publication Date]) AND
"supplementary"[Filter]
~~~

Datasets in the Hemberg collection (https://hemberg-lab.github.io/scRNA.seq.datasets/) were merged into this list. Only animal single-cell transcriptomic datasets profiling samples of normal conditions were selected. We also manually filtered small-scale or low-quality data. Additionally, several other high-quality datasets missing in the previous list were included for comprehensiveness.

The expression matrices and metadata of selected datasets were retrieved from GEO, supplementary files of the publication or by directly contacting the authors. Metadata were further manually curated by adding additional descriptions in the paper to acquire the most detailed information of each cell. We unified raw cell type annotation by Cell Ontology^42^, a structured controlled vocabulary for cell types. Closest Cell Ontology terms were manually assigned based on the Cell Ontology description and context of the study.

### Building reference panels for the ACA database

Two types of searchable reference panels are built for the ACA database. The first consists of individual datasets with dedicated models trained on each, while the second consists of datasets grouped by organ and species, with models trained to align multiple datasets profiling the same species and same organ.

Data preprocessing follows the same procedure as in previous benchmarks. Both cross-dataset batch effect and within-dataset batch effect are manually examined and removed when necessary. For the first type of reference panels, datasets too small (typically < 1,000 cells sequenced) are excluded because of insufficient training data. These datasets are still included in the second type of panels, where they are trained jointly with other datasets profiling the same organ in the same species. For each reference panel, four models with different starting points are trained.

### Web interface

For conveniently performing and visualizing Cell BLAST analysis, we built a one-stop Web interface. The client-side was made from Vue.js, a single-page application Javascript framework, and D3.js for cell ontology visualization. We used Koa2, a web framework for Node.js, as the server side. The Cell BLAST Web portal with all accessible curated datasets is deployed on Huawei Cloud.

---

**Supplementary Fig. 1.**
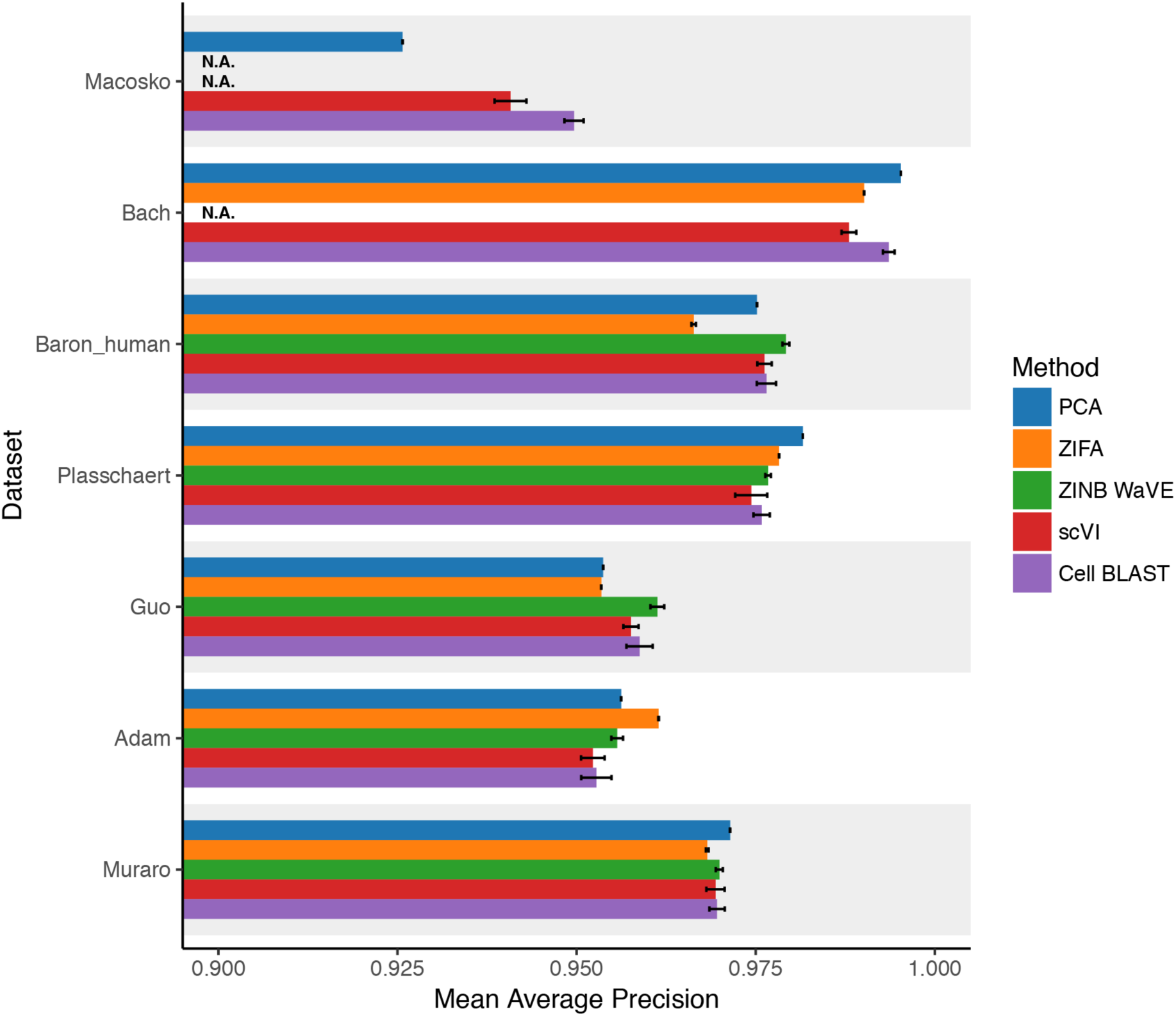
Comparing low-dimensional space cell type resolutions of different dimension reduction methods. Nearest neighbor cell type mean average precision (MAP) is used to evaluate how well biological similarity is captured. MAP can be thought of as a generalization to nearest neighbor accuracy, with larger values indicating higher cell type resolution and, thus, more suitable for cell querying. Error bars indicate mean ± s.d. Methods that did not finish under the 2-hour time limit are marked as N.A.

**Supplementary Fig. 2.**
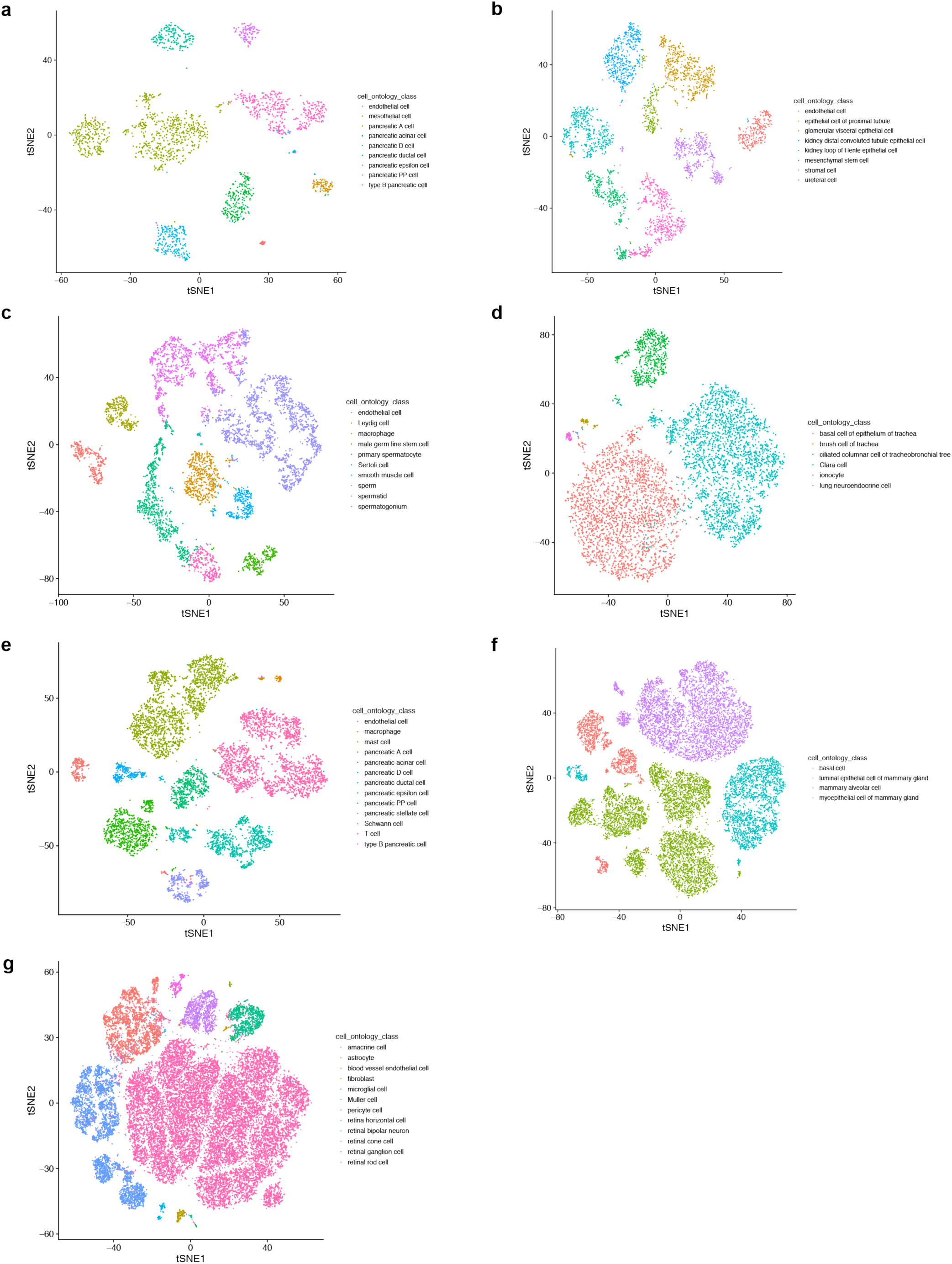
t-SNE visualization of latent spaces learned by our model. (**a**) “Muraro”^43^, (**b**) “Adam”^44^, (**c**) “Guo”^45^, (**d**) “Plasschaert”^15^, (**e**) “Baron_human”^46^, (**f**) “Bach”^47^, (**g**) “Macosko”^48^.

**Supplementary Fig. 3.**
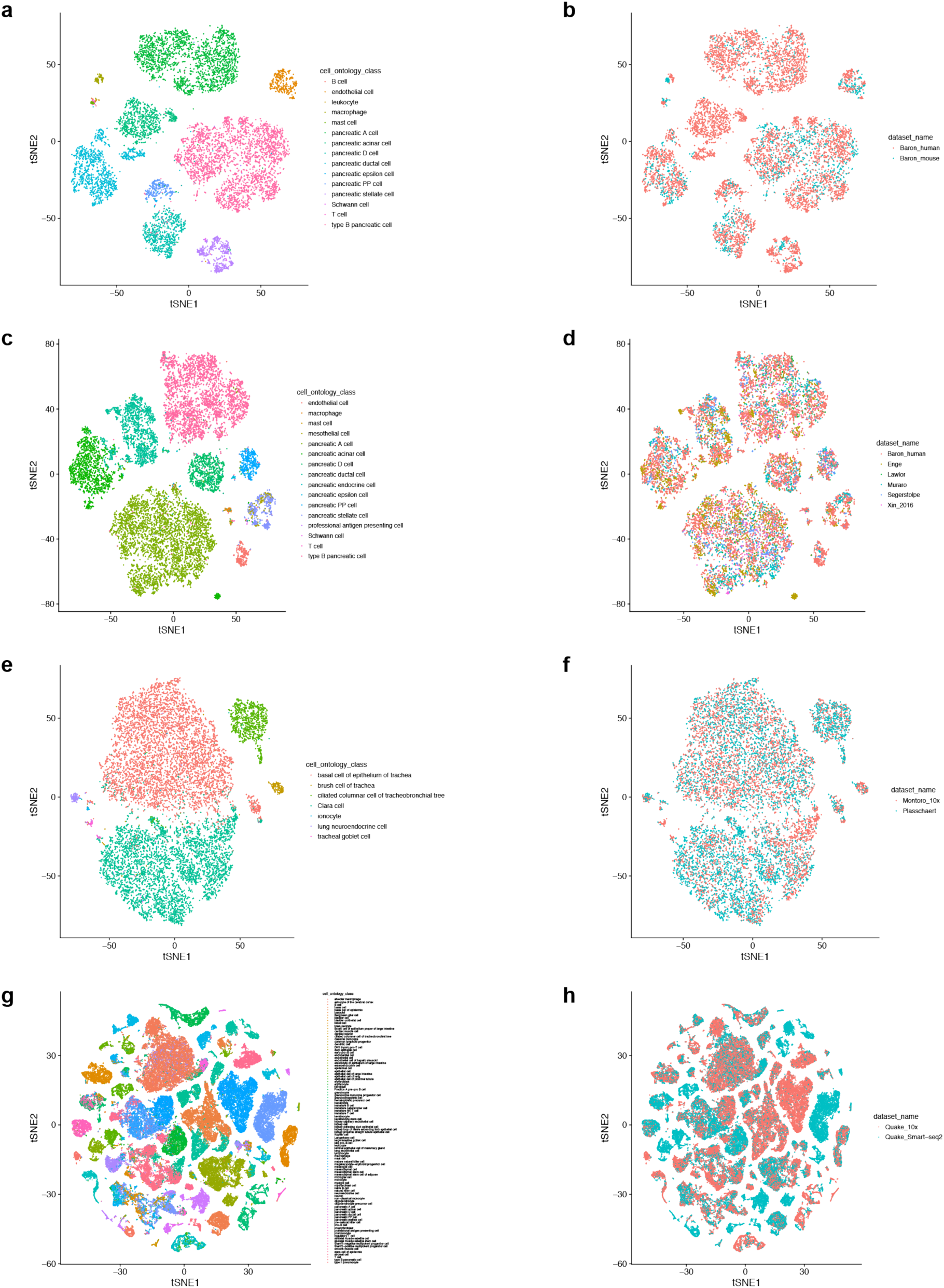
t-SNE visualization of latent spaces learned by our model on combinations of multiple datasets, with batch effect corrected. Figures in the left column color cells by cell type, while figures in the right column color cells by dataset. (**a**-**b**) “Baron_human”^46^ and “Baron_mouse”^46^; (**c**-**d**) “Baron_human”^46^, “Muraro”^43^, “Enge”^49^, “Segerstolpe”^50^, “Xin_2016”^51^ and “Lawlor”^52^; (**e**-**f**) “Montoro_10x”^14^ and “Plasschaert”^15^; (**g**-**h**) “Quake_Smart-seq2”^18^ and “Quake_10x”^18^.

**Supplementary Fig. 4.**
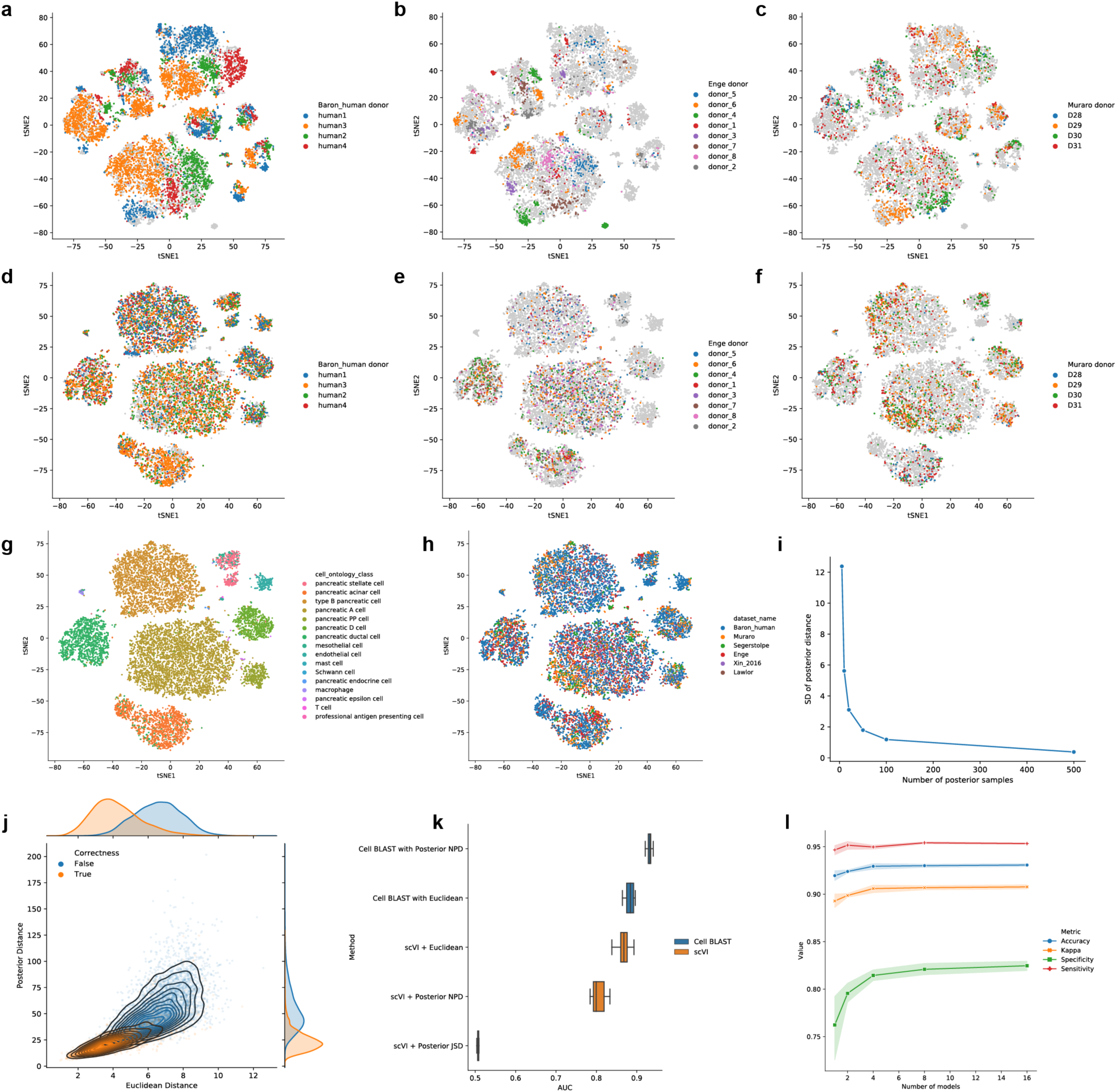
Multilevel batch effect correction and Cell BLAST strategy optimization. (**a**-**c**) Latent space learned with only cross-dataset batch effect correction, colored by (**a**) donor in “Baron_human”^46^, (**b**) donor in “Enge”^49^, (**c**) donor in “Muraro”^43^. (**d**-**h**) Latent space learned with both cross-dataset and within-dataset batch effect correction, colored by (**d**) donor in “Baron_human”^46^, (**e**) donor in “Enge”^49^, (**f**) donor in “Muraro”^43^, (**g**) cell type, (**h**) dataset. (**i**) Standard deviation decreases as the number of samples from the posterior increases. (**j**) Relationship between Euclidean distance and NPD in “Baron_human”^46^ data. The orange points represent cell pairs that are of the same cell type (“positive pairs”), while the blue points represent cell pairs of different cell types (“negative pairs”). (**k**) AUROC of different distance metrics in discriminating cell pairs with the same cell type from cell pairs with different cell types. Box plots indicate the median (center lines), interquantile range (hinges), and 1.5 times the interquantile range (whiskers). Note that the posterior distribution distances for scVI only lead to a decrease in performance, possibly due to improper Gaussian assumption in the posterior. (**l**) Accuracy, Cohen’s κ (a measure of prediction accuracy corrected for chance, see **Methods** for more details)^2^, specificity and sensitivity all increase as the number of models used for cell querying increases, among which the improvement of specificity is the most significant. Error bars indicate mean ± s.d.

**Supplementary Fig. 5.**
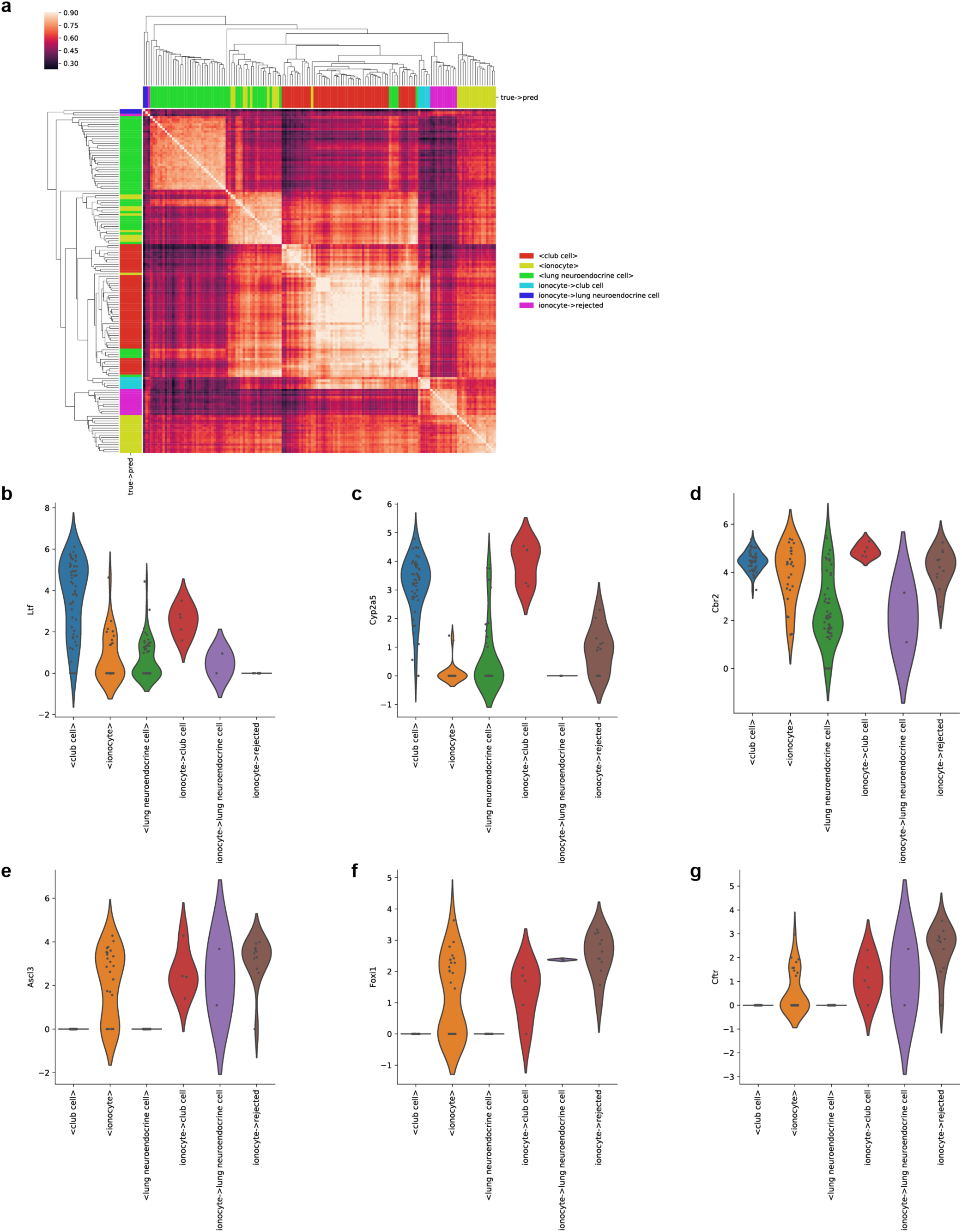
Ionocytes predicted to be club cells are potentially doublets or of an intermediate cell state. (**a**) Cell-cell correlation heatmap for several cell types of interest. Cells labeled as “<X>” are reference cells in the “Montoro”^14^ dataset. Cells labeled as “X->Y” are cells annotated as “X” in the original “Plasschaert”^15^ dataset but predicted to be “Y”. (**b**-**d**) Expression levels of several club cell markers in the cell groups of interest. (**e**-**g**) Expression levels of several ionocyte markers in the cell groups of interest.

**Supplementary Fig. 6.**
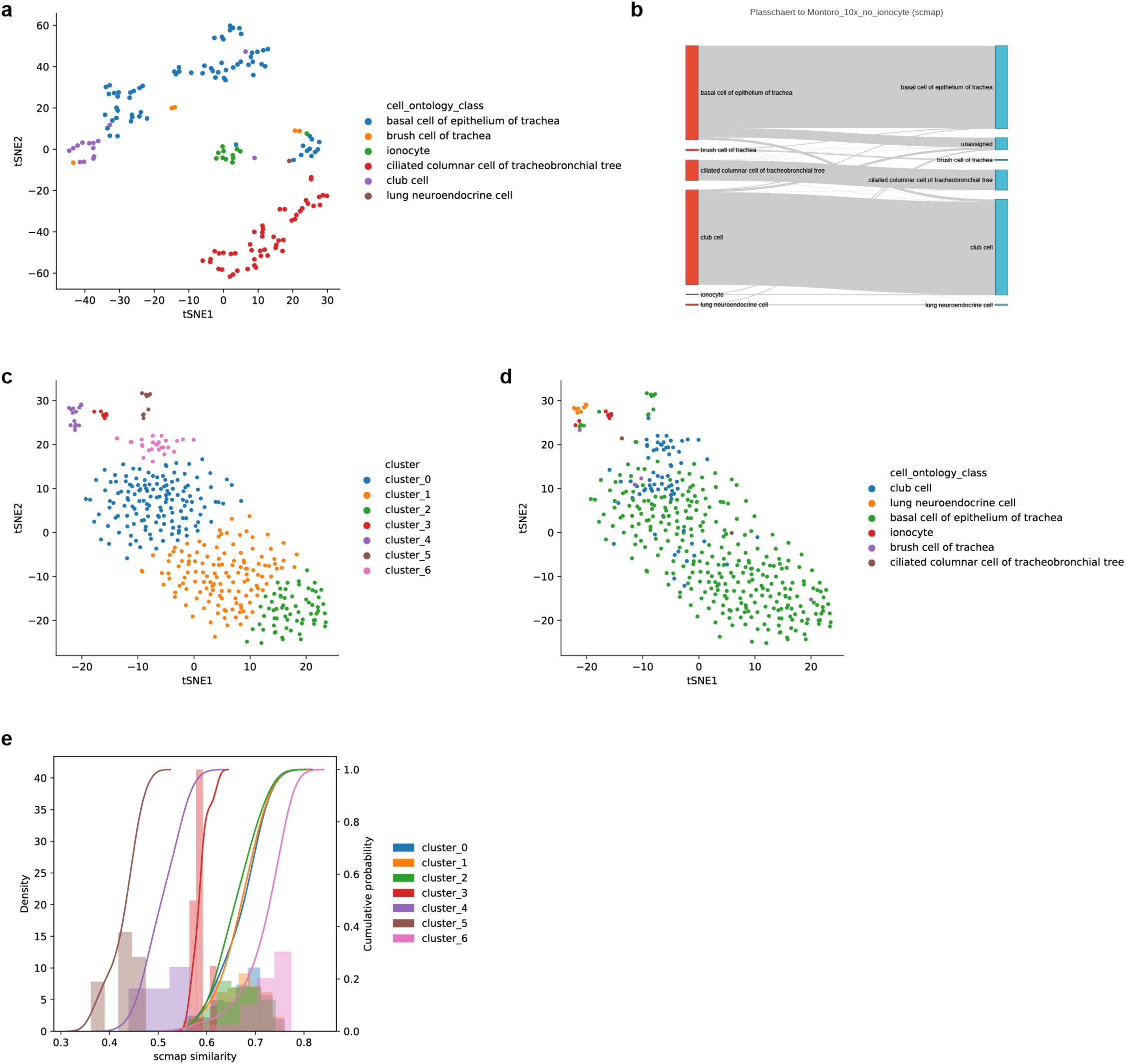
Rejected cells in the “Montoro” - “Plasschaert” query. (**a**) t-SNE visualization of Cell BLAST-rejected cells, colored by cell type. (**b**) Sankey plot of scmap prediction. (**c**, **d**) t-SNE visualization of scmap-rejected cells, colored by unsupervised clustering (**c**) and cell type (**d**). (**e**) scmap similarity distribution in each cluster of scmap-rejected cells. The rejected ionocytes do not have the lowest cosine similarity scores to draw sufficient attention.

**Supplementary Fig. 7.**
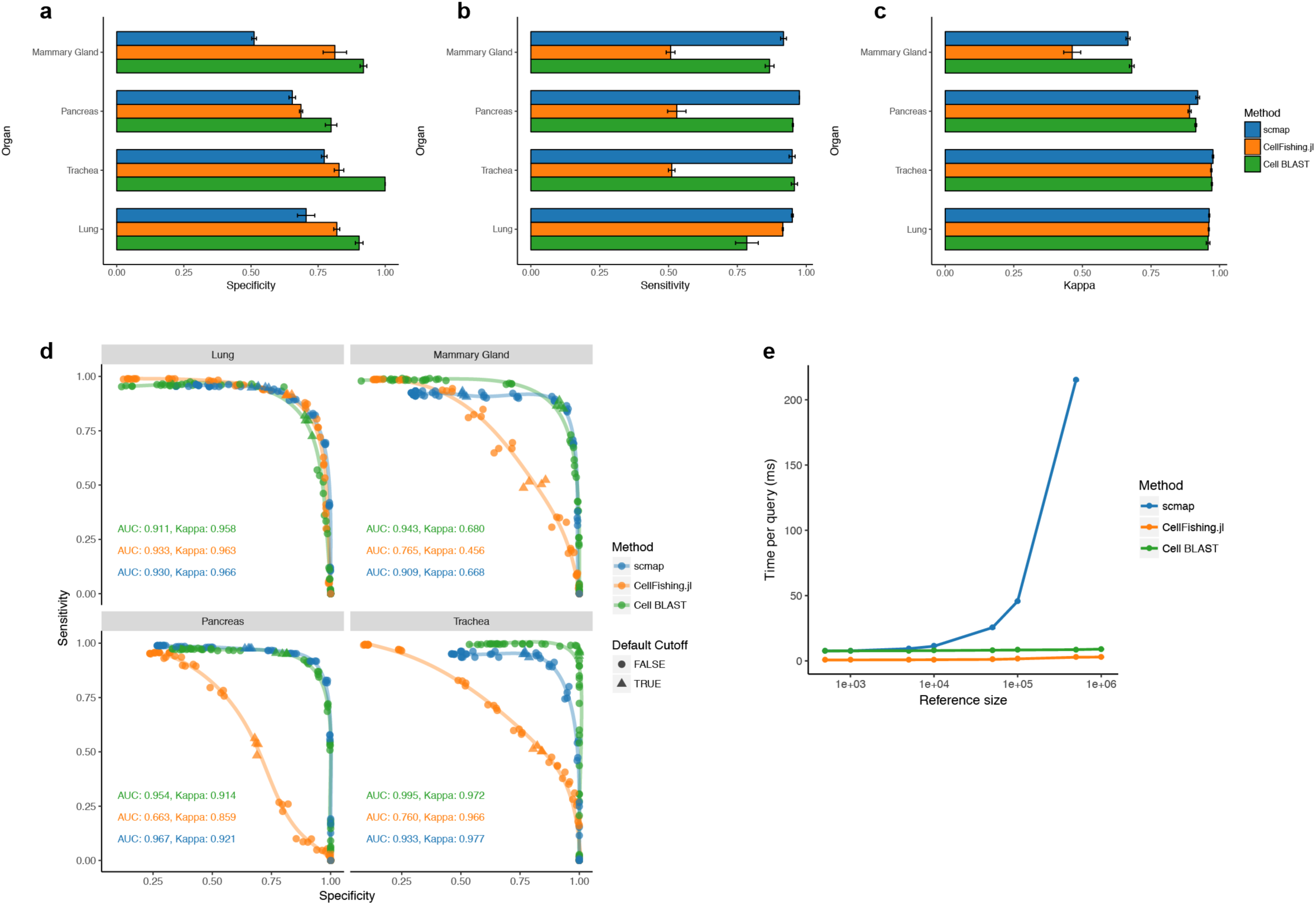
Benchmarking query-based cell typing. (**a**-**c**) Querying specificity (**a**), sensitivity (**b**) and Cohen’s κ (**c**) for different methods under the default setting. Error bars indicate mean ± s.d. (**d**) ROC curve of cell querying in four different groups of test datasets. Cohen’s κ values in the bottom left of each subpanel correspond to the optimal point on the ROC curve. Points corresponding to each method’s default cutoff (scmap: cosine distance = 0.5, CellFishing.jl: Hamming distance = 110, Cell BLAST: p-value = 0.05) are marked as triangles. Note that CellFishing.jl does not provide a default cutoff, so we chose a Hamming distance of 110, which is the closest to balancing sensitivity and specificity, but it is still far from being stable across different datasets. (**e**) Querying speed on reference datasets of different sizes subsampled from the 1M mouse brain dataset^35^. Error bars indicate mean ± s.d.

**Supplementary Fig. 8.**
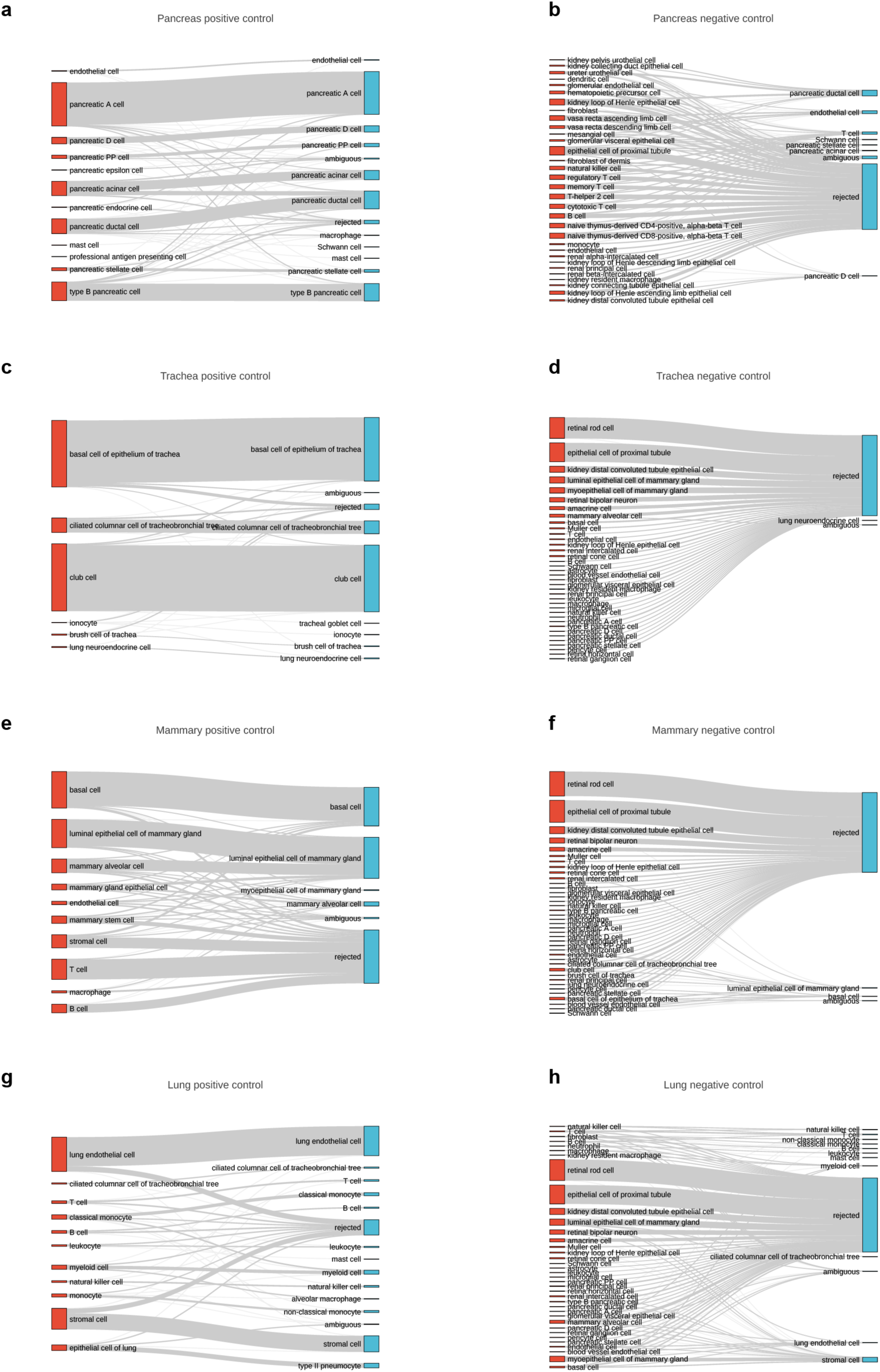
Sankey plots for Cell BLAST in the cell-querying benchmark.

**Supplementary Fig. 9.**
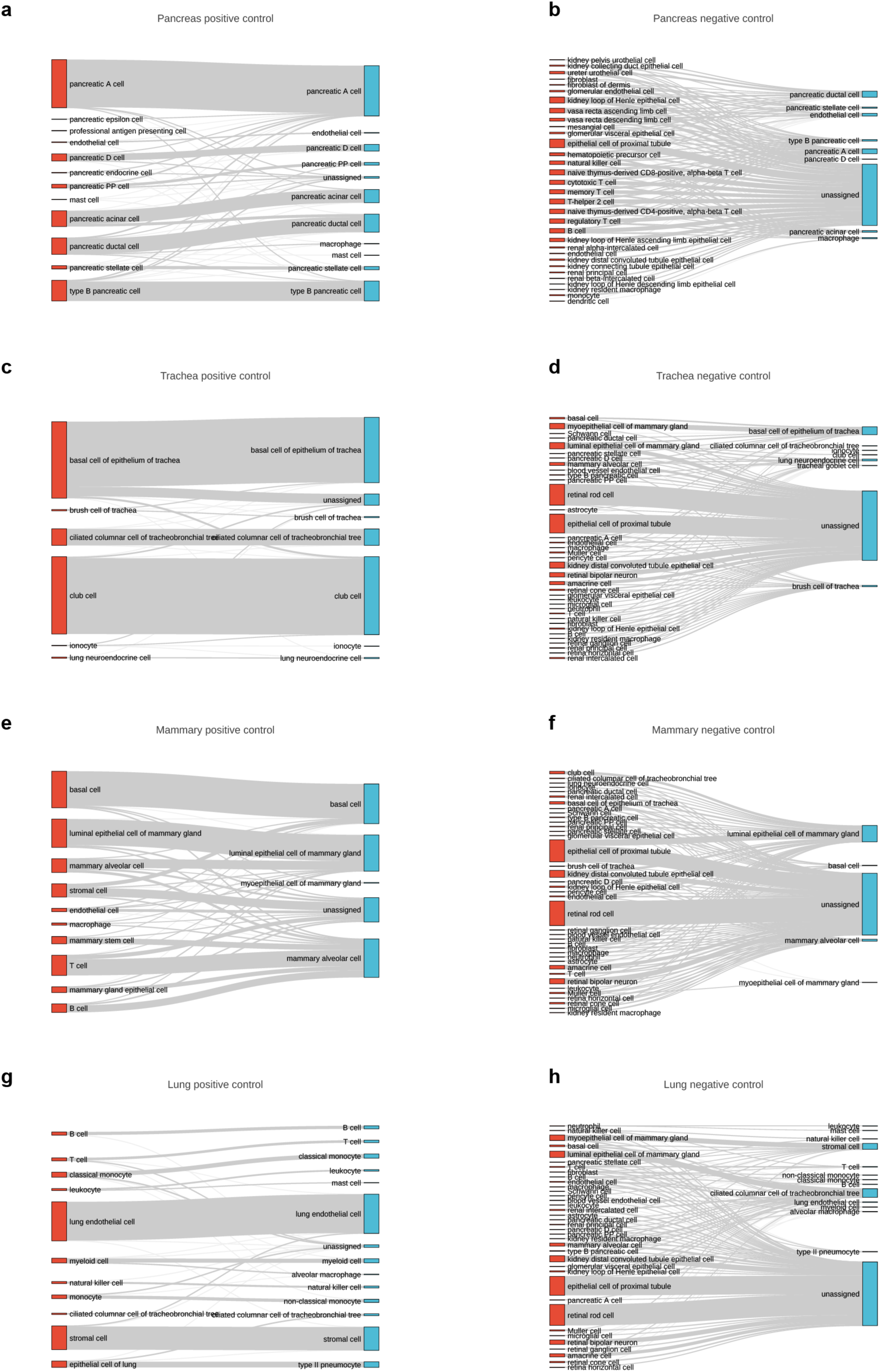
Sankey plots for scmap in the cell-querying benchmark.

**Supplementary Fig. 10.**
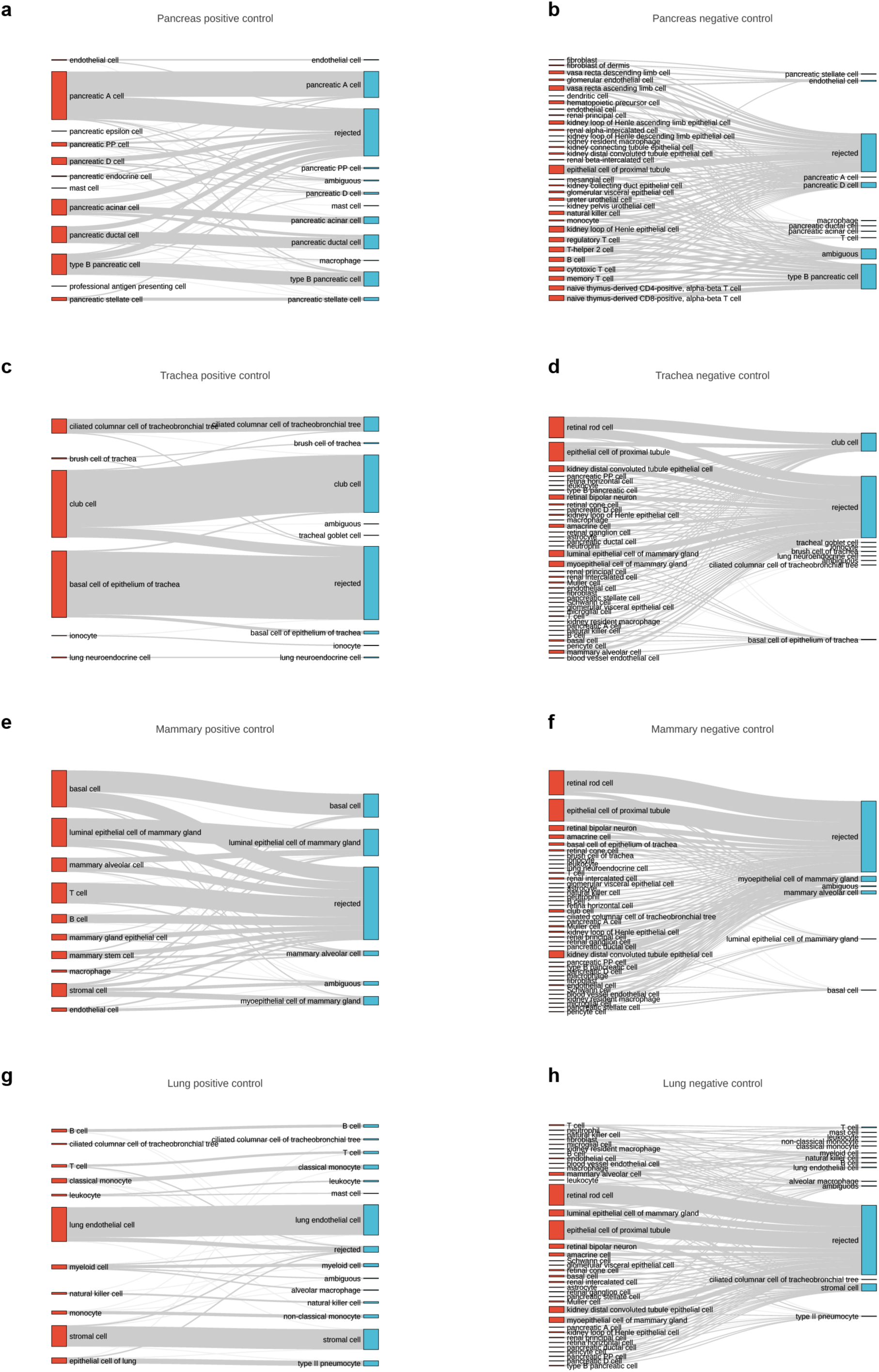
Sankey plots for CellFishing.jl in the cell-querying benchmark.

**Supplementary Fig. 11.**
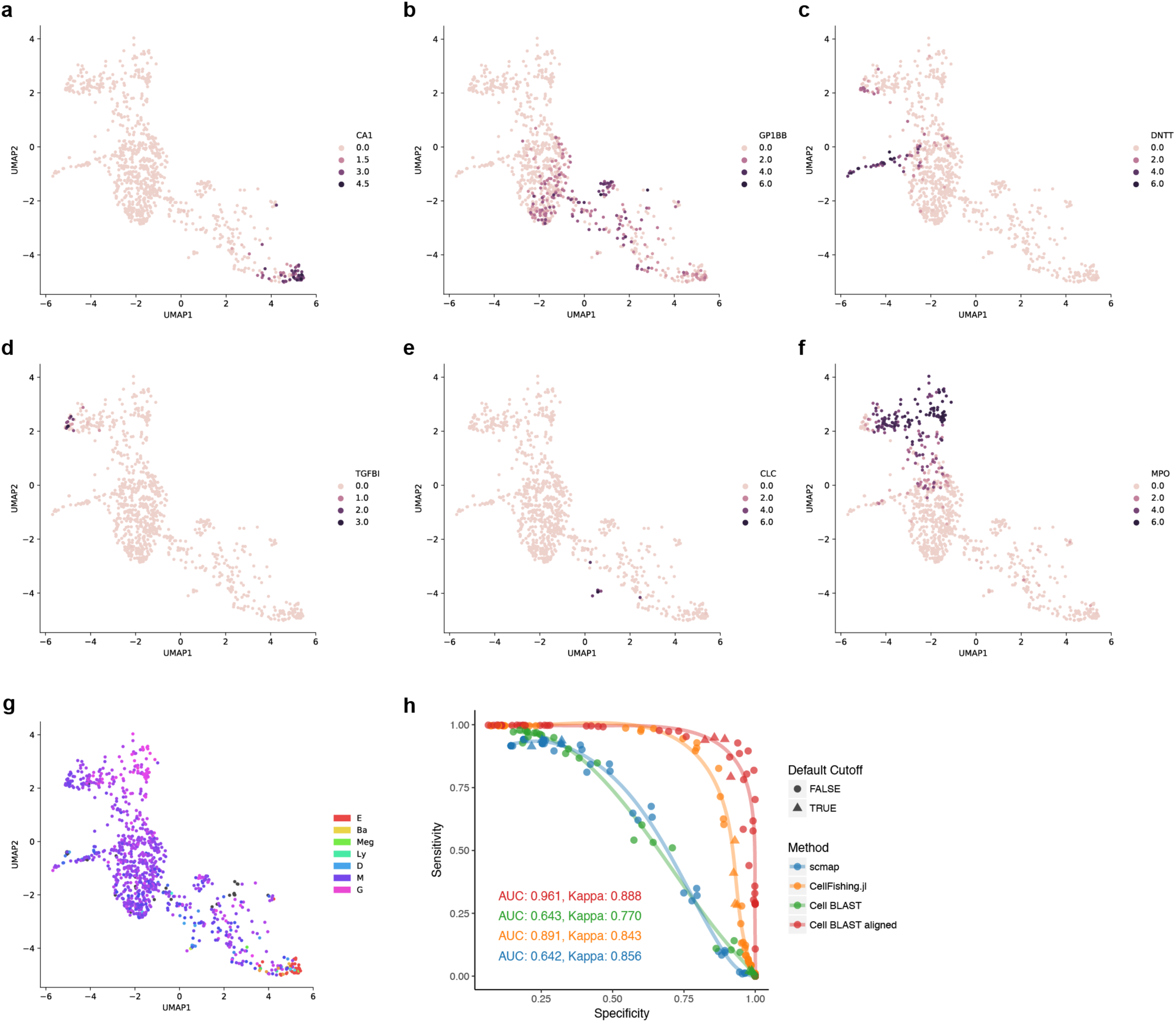
Using “online tuning” in hematopoietic progenitor and *Tabula Muris*^18^ spleen data. UMAP visualization of the “Velten”^17^ dataset, colored by the expression of known lineage markers, including CA1 for the erythrocyte lineage (**a**), GP1BB for the megakaryocyte lineage (**b**), DNTT for the B-cell lineage (**c**), TGFBI for monocyte and dendritic cell lineages (**d**), CLC for eosinophil, basophil, and mast cell lineages (**e**), MPO for the neutrophil lineage (**f**), and scmap predicted cell fate distribution (**g**). (**h**) ROC curve of cell querying in *Tabula Muris*^18^ spleen data. Cohen’s κ values in the bottom left of each subpanel correspond to the optimal point on the ROC curve. Points corresponding to each method’s default cutoff (scmap: cosine distance = 0.5, CellFishing.jl: Hamming distance = 110, Cell BLAST: p-value = 0.05) are marked as triangles.

**Supplementary Fig. 12.**
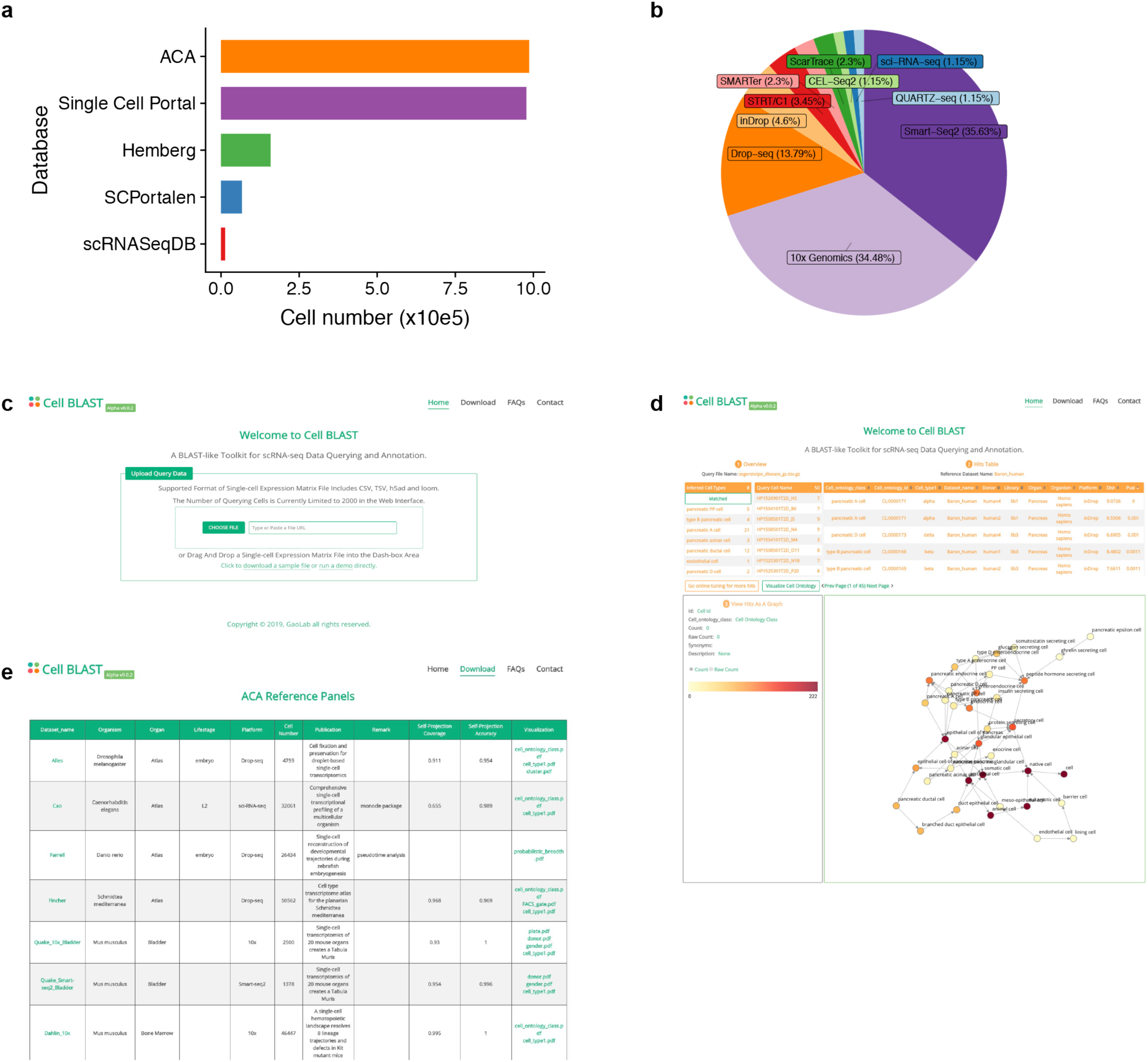
ACA database and Cell BLAST Web portal. (**a**) Comparison of cell numbers in different single-cell transcriptomics databases. (**b**) Composition of different single-cell sequencing platforms in ACA. (**c**) Home page of the Cell BLAST Web interface. (**d**) Web interface showing the results of a sample query. (**e**) A full list of ACA reference panels available in our Web interface.

**Supplementary Table 1.**
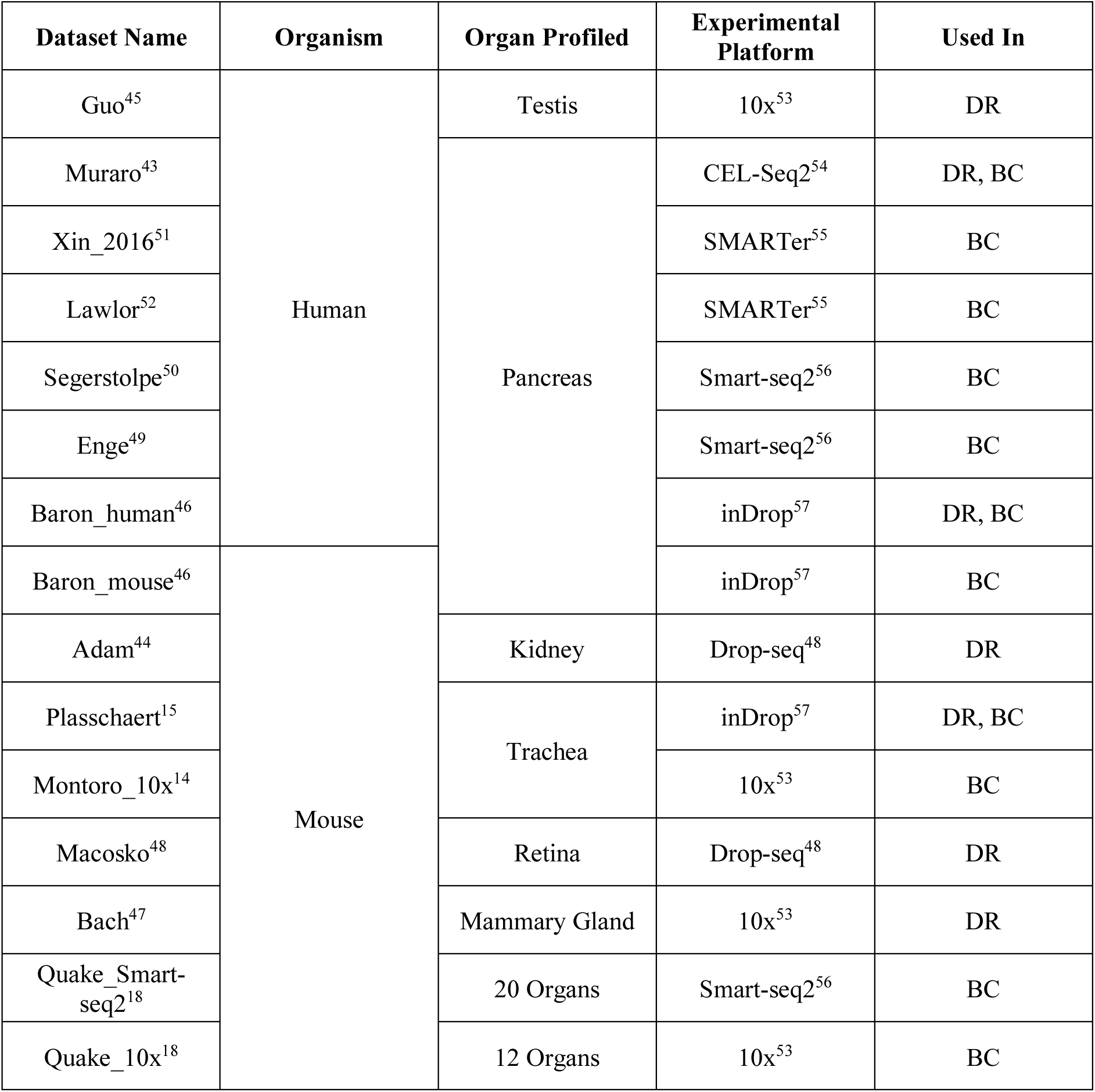
Datasets used in dimensionality reduction and batch effect correction benchmarking. DR, dimension reduction benchmarking; BC, batch effect correction benchmarking.

**Supplementary Table 2.**
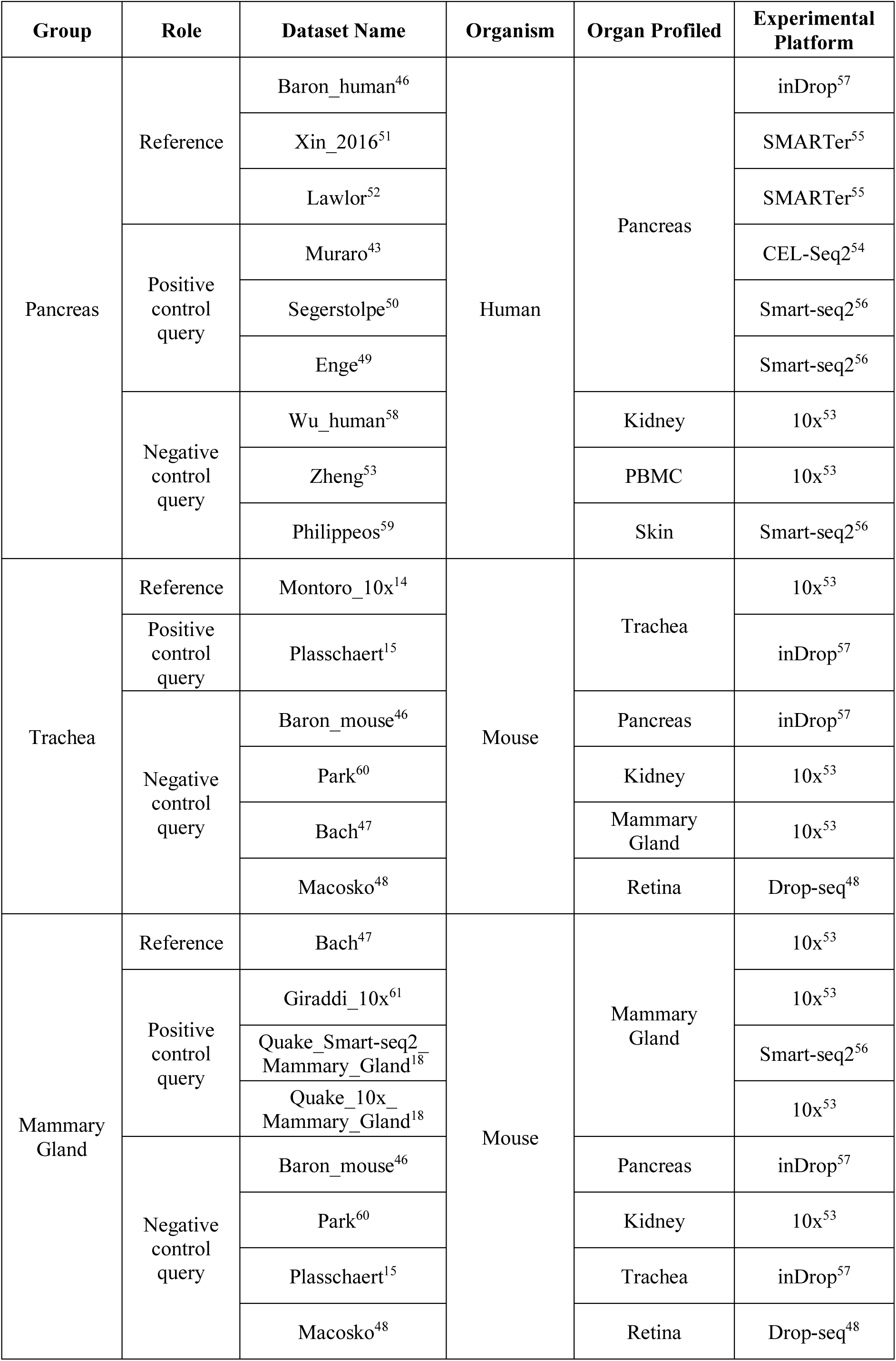
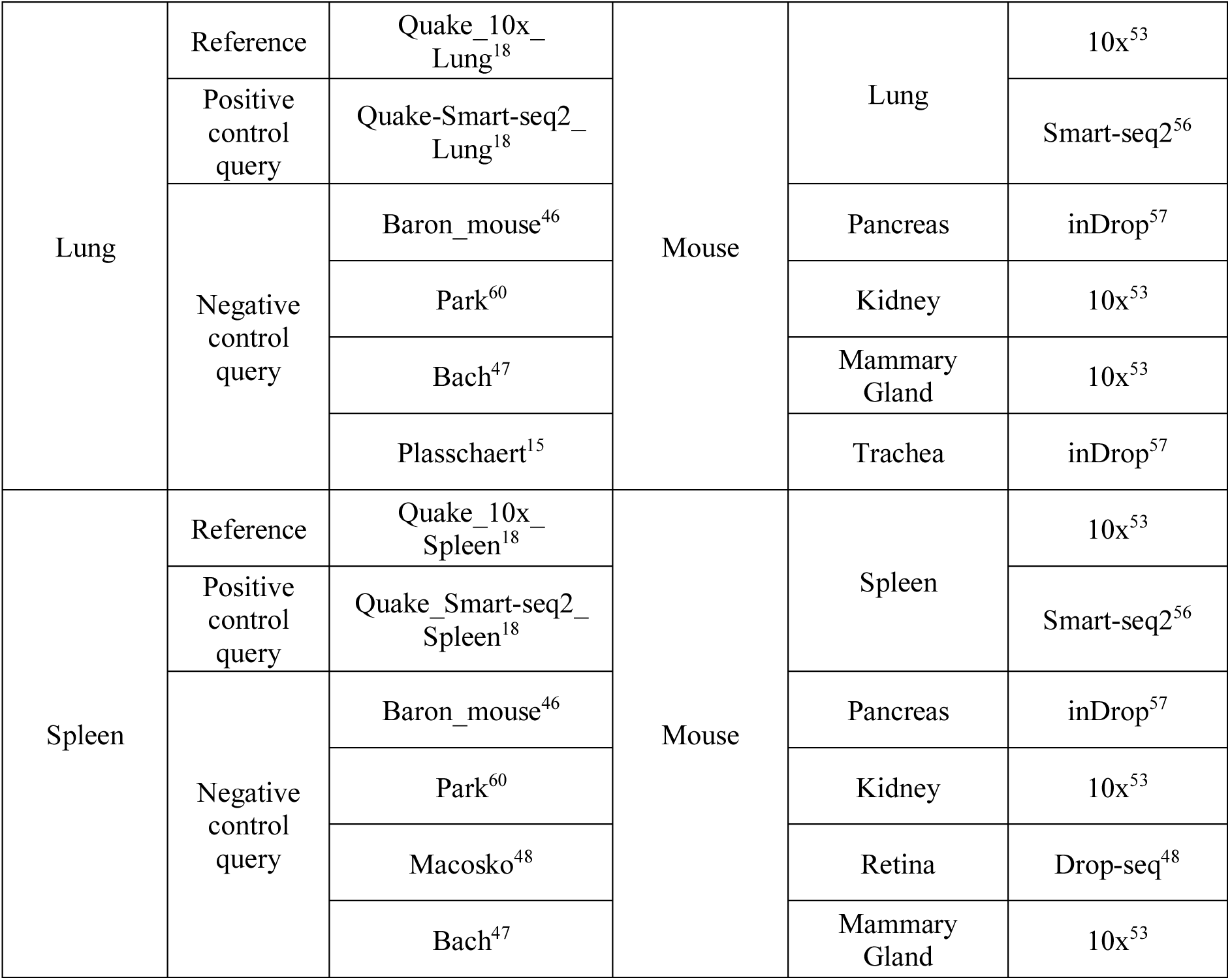
Datasets used in cell query benchmarking.

**Supplementary Table 3. Raw data of benchmarking results.**

**Supplementary Table 4. Datasets in ACA.**

